# TNFR2 costimulation differentially impacts regulatory and conventional CD4^+^ T-cell metabolism

**DOI:** 10.1101/2022.03.25.485786

**Authors:** Mark Mensink, Esther A. Zaal, Thi Tran Ngoc Minh, Ellen Schrama, Celia R. Berkers, Jannie Borst, Sander de Kivit

## Abstract

CD4^+^ conventional T cells (Tconvs) mediate adaptive immune responses, whereas regulatory T cells (Tregs) suppress those responses to safeguard the body from autoimmunity and inflammatory diseases. The opposing activities of Tconvs and Tregs depend on the stage of the immune response and their environment, with an orchestrating role for cytokine- and costimulatory receptors. Nutrient availability also impacts T-cell functionality via metabolic and biosynthetic processes that are largely unexplored. Many data argue that costimulation by Tumor Necrosis Factor Receptor 2 (TNFR2) favors support of Treg over Tconv responses and therefore TNFR2 is a key clinical target. Here, we review the pertinent literature on this topic and highlight the newly identified role of TNFR2 as a metabolic regulator for thymus-derived (t)Tregs. We present novel transcriptomic and metabolomic data that show the differential impact of TNFR2 on Tconv and tTreg gene expression and reveal distinct metabolic impact on both cell types.

## 1 Introduction

Adaptive immunity is controlled by the opposing activities of Tconvs and Tregs. The Treg lineage is hallmarked by expression of the master transcriptional regulator FOXP3 that, in conjunction with other transcription factors, dictates Treg function (1, 2). CD4^+^ T cells may gain FOXP3 expression during thymic development and thus become tTregs that have a T-cell receptor (TCR) repertoire focused on self-antigens. Alternatively, CD4^+^ Tconvs may gain FOXP3 expression and convert into peripherally induced (p)Tregs during responses to foreign antigens (1, 2). In most tissues, specialized tTreg subsets reside that respond and adapt to Tconv responses (3). Tregs prevent autoimmunity, maintain tissue homeostasis and limit inflammation. Aberrant Treg function may therefore contribute to immune disorders or cancer (4-6). To improve immunotherapy of such diseases, we must understand the signals that control Tconv and Treg responses. These responses are initiated in lymphoid organs, but continue and often persist in many non-lymphoid tissues in health and disease (3). Both T-cell types are activated by engagement of their TCR, which together with costimulatory- and cytokine receptor stimulation leads to proliferation and effector differentiation (7). Besides these signals that are primarily delivered by dendritic cells (DCs), other environmental signals such as nutrient availability impact T-cell responses (8). Especially in rapidly proliferating cells, nutrients provide the metabolic building blocks for biosynthetic pathways (9). Therefore, it can be envisioned that the interplay of immune receptors and nutrient metabolism is important for the activity of Tconvs and Tregs.

Tconvs and Tregs share numerous immunoregulatory receptors. These may differentially impact the response of each cell type, due to differential wiring of signal transduction pathways (10) and other physiological differences between the cell types. Accumulating data emphasize the selective importance of the T-cell costimulatory receptor TNFR2 for promoting Treg as opposed to Tconv responses. In mouse models, TNFR2-stimulated Treg responses were shown to protect from autoimmunity and graft-versus-host disease (11-15). We recently reported that TNFR2 costimulation causes a glycolytic switch in activated human tTregs, providing the first evidence that TNFR2 can regulate cell metabolism (16). Given the great interest in antibody-based TNFR2 targeting in immune-related diseases, it is important to understand the consequences of TNFR2 costimulation for Tconv and Treg responses. At present, data suggest that TNFR2 primarily promotes Treg responses, while leaving Tconvs unaffected. Here, we review the validity of this assumption and present recent data from our own laboratory that highlight the response of tTregs and Tconvs to TNFR2 costimulation at the metabolic level. TNFR2 differentially regulates metabolism of tTregs and Tconvs, which may have implications for physiology, cell therapy and drug targeting.

## 2 Rationale for TNFR2 targeting in immune-related diseases

### 2.1 TNF(R) targeting in disease

Tumor Necrosis Factor (TNF) is a cytokine involved in a plethora of biological processes and is well known for its pro-inflammatory properties (17). Its involvement in the pathogenesis of autoimmune and chronic inflammatory diseases such as rheumatoid arthritis has led to the development and broad clinical application of TNF-blocking agents, including the antibodies infliximab, adalimumab and golimumab and the soluble IgG1-TNFR2 fusion protein etanercept (18, 19). Despite the successes of TNF-blocking therapies, these treatments are ineffective for part of the patients, can have side effects and can even exacerbate the disease (20-24). This calls for a better understanding of the mechanistic consequences of TNF inhibition.

TNF is initially expressed on the cell surface as a transmembrane molecule and cleaved to be released in its soluble form (25-27). Either soluble or transmembrane TNF can activate TNFR1 (TNFRSF1A), while transmembrane TNF primarily activates TNFR2 (TNFRSF1B) (28, 29). TNFR1 and TNFR2 have different expression patterns, distinct intracellular domains and evoke different cellular responses (30). TNFR1 is a ubiquitously expressed receptor (https://www.proteinatlas.org/) that can mediate inflammation and different forms of cell death (17). TNFR2 has a more restricted tissue distribution, is more prone to anti-inflammatory signaling and mediates cell survival and proliferation (31).

It is already known for decades that TNF can have both pro-inflammatory and anti-inflammatory effects (32-34). Neutralization of TNF in immune-related diseases aims to block the pro-inflammatory effects of TNF–TNFR1 signaling (35). However, such intervention will also block the anti-inflammatory properties of TNF, which seem to largely result from TNFR2 signaling into Tregs (36). Thus, drugs interfering with TNF function may restrain Treg responses and therefore be pro-instead of anti-inflammatory. Drugs that specifically modulate either TNFR1 or TNFR2 activity will therefore be more suited to combat inflammatory and autoimmune diseases and transplant rejection, as has been reviewed comprehensively (31, 35, 37-41).

### 2.2 The role of TNFR2 on Tregs and Tconvs

TNFR2 is highly expressed by a fraction of human and mouse Tregs, on which it supports FOXP3 expression and proliferation and maintains suppressive activity (11, 16, 42-46). Specific polymorphisms in the human *TNFRSF1B* gene leading to loss of TNFR2 expression or altered function are associated with inflammatory bowel disease, ankylosing spondylitis, lupus and/or rheumatoid arthritis, supporting the idea that TNFR2 protects against these diseases (31, 47-53). In mouse models, further evidence has been gathered that support this notion. Wild-type but not TNFR2-deficient Tregs could inhibit experimental colitis in mice (11). In experimental autoimmune encephalomyelitis (EAE), a mouse model for multiple sclerosis, Treg-restricted TNFR2 deficiency led to a reduction in expression of Treg signature molecules and suppressive function, which was associated with disease exacerbation (12). Another study supports the notion that Tregs require TNFR2 expression to suppress EAE (13). Tregs also selectively expanded upon TNFR2 agonism and reduced graft-versus-host disease in a TNFR2-dependent manner in allogeneic hematopoietic stem cell transplantation in mice (14, 15). Whereas promoting TNFR2 activity on Tregs could be promising in autoimmune diseases and transplantation, TNFR2 inhibition is considered in cancer therapy (54). High TNFR2 expression is observed on certain cancers and on tumor-associated Tregs (54-58). TNFR2 agonism and antagonism are currently being explored in both autoimmune diseases and cancer, potentially opening new treatment avenues (57, 59-61).

TNFR2 functions as a costimulatory receptor not only on Tregs, but also on Tconvs by lowering the threshold of T-cell activation and increasing survival and proliferation (62-65). TNFR2 expression is upregulated on Tconvs upon TCR activation and is associated with resistance to Treg-mediated suppression in mice, as determined by increased proliferation and effector cytokine production (42, 66). TNFR2-deficient Tconvs failed to induce colitis in mice, which was linked to reduced proliferation and less IFN-γ production (67). Together, these findings indicate that TNFR2 has costimulatory effects on both Tconvs and Tregs, but that under specific conditions, Treg responsiveness prevails. In this Perspective, we offer our recent experimental findings resulting from unbiased “omics” approaches that give new insights into the effects of TNFR2 agonism on both tTregs and Tconvs. We believe that this information can stimulate new research directions with clinical relevance, in particular regarding the impact of environmental metabolic cues on the Treg/Tconv balance and functional outcomes.

We present an unbiased, side-by-side analysis of the consequences of TNFR2 costimulation on human Tconvs and tTregs at the level of gene expression and cell metabolism. The Treg lineage consists of tTregs and pTregs (68, 69), but pTregs are largely restricted to mucosal surfaces and the maternal-fetal interface (70, 71). The majority of Tregs in vivo are tTregs, which control systemic and tissue-specific autoimmunity (3, 72). Therefore, we focused on tTregs that were isolated to high purity from blood as described before (16, 73).

## 3 Effects of TNFR2 on Tregs and Tconvs according to our recent work

### 3.1 Transcriptomics identifies responses of Tconvs and tTregs to TNFR2 costimulation

Contrary to Tconvs, Tregs rely on costimulation to initiate proliferation after in vitro triggering of the TCR/CD3 complex by anti-CD3 antibody (74). In addition, Tregs require exogenous IL-2 in such assays, since they cannot produce this cytokine after activation, unlike Tconvs (75, 76). Accordingly, in our experimental setting, Tconvs already started to proliferate upon stimulation of the TCR/CD3 complex alone, whereas tTregs required additional costimulation by either CD28 or TNFR2 (**Figure 1A**) (16). We used a comparative setting of Tconvs and tTregs that had been pre-expanded, rested and restimulated via CD3 alone, CD3/CD28 or CD3/TNFR2. After 24 h, cells were harvested for transcriptome analysis (16).

**Figure 1.**
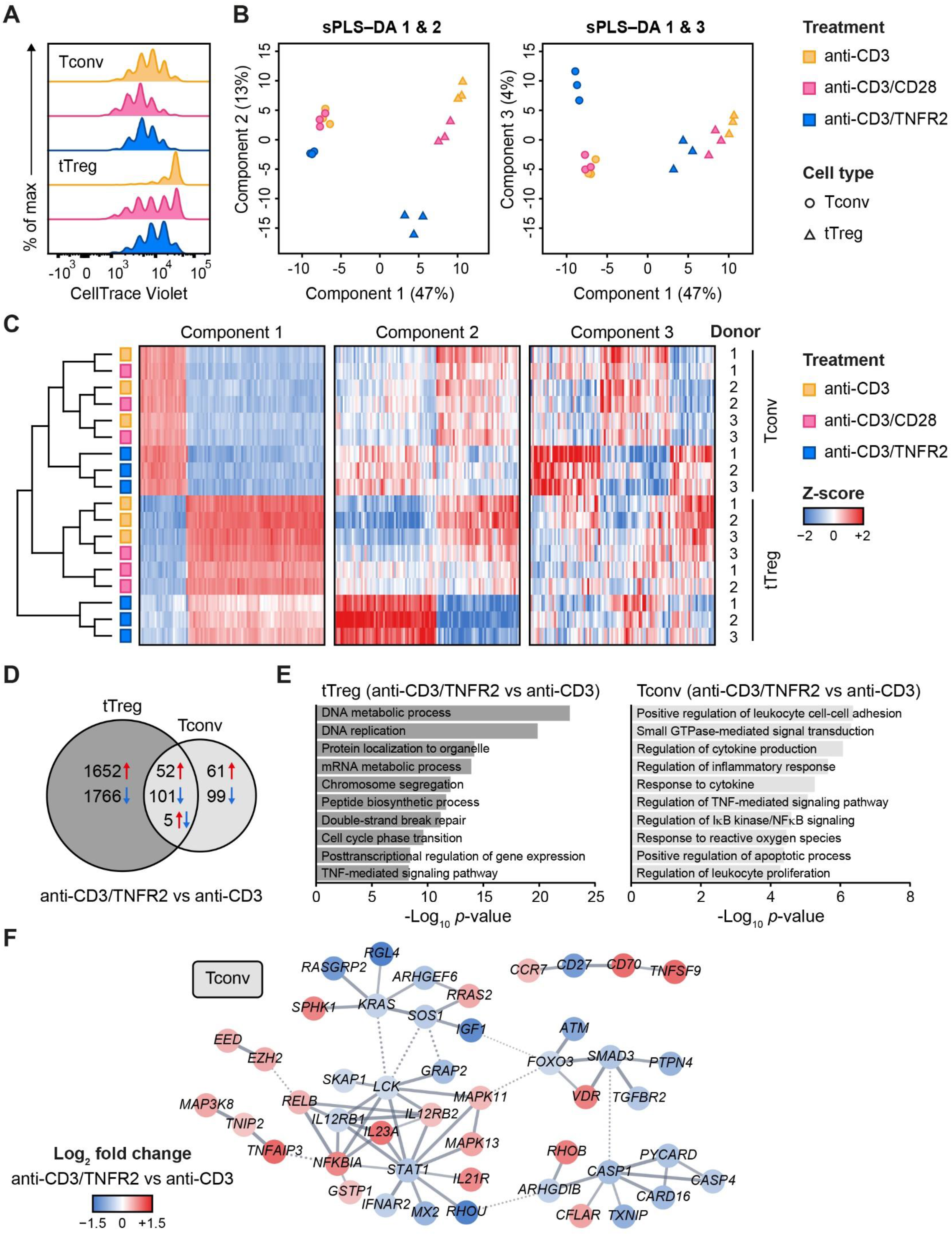
Transcriptomic analysis of human Tconvs and tTregs highlights distinct responses upon CD3, CD3/CD28 or CD3/TNFR2 stimulation. Methods for cell isolation, cell culture and RNA sequencing were described before (16). In brief, naïve Tconvs and tTregs were isolated from healthy donor blood based on CD4^+^CD25^lo^CD127^hi^CD45RA^+^GPA33^int^ and CD4^+^CD25^hi^CD127^lo^CD45RA^+^GPA33^hi^ phenotypes, respectively. Cells were cultured for 2 weeks in IMDM (Gibco, Life Technologies), supplemented with 8% FCS (Sigma) and penicillin/streptomycin (Roche), in presence of IL-2 (DuPont Medical) and agonistic mAbs against CD3 (clone CLB-T3/4.E, IgE, Sanquin) and CD28 (clone CLB-CD28/1, Sanquin). Following expansion, cells were cultured for 4 days in the absence of anti-CD3 and anti-CD28 mAbs. Next, Tconvs and tTregs were restimulated for 24 h in presence of IL-2 and either anti-CD3 mAb alone or combined with anti-CD28 or anti-TNFR2 (clone MR2-1, Hycult Biotech) mAb, and subjected to transcriptomics (*n* = 3). Using Qlucore Omics Explorer (version 3.8), we performed trimmed mean of M values (TMM) normalization (109) and only included transcripts with non-zero read counts for all samples. Next, we performed sPLS–DA using R (version 4.1.0) by applying the *splsda* function embedded in the mixOmics package (version 6.16.3) (77, 110). **(A)** After resting, pre-expanded Tconvs and tTregs were labeled with CellTrace Violet (Invitrogen) and stimulated for 96 h to assess cell proliferation by flow cytometry. For color legend of stimuli, see panel **B**. **(B)** Clustering of indicated sample groups by components 1–3 using sPLS–DA. **(C)** Representative heat map of the sPLS–DA results, showing hierarchical clustering of indicated sample groups and relative levels (z-scores) of the top 100 transcripts per component 1–3. **(D)** Venn diagram depicting the number of up- (red) and downregulated (blue) transcripts in CD3/TNFR2-versus CD3-stimulated samples, including unique and shared changes in tTregs and Tconvs (*p* < 0.05, log_2_ fold change > 0.32 or < −0.32). **(E)** GO Biological Process enrichment analysis of the differentially expressed genes shown in **D**. **(F)** STRING (version 11.5) network of genes involved in the enriched processes for Tconvs in **E**, including only genes with high-confidence associations. Markov Cluster Algorithm (MCL) clustering (111) was performed and inter-cluster edges are displayed as dotted lines. Genes were colored in Cytoscape (version 3.9.0) based on log_2_ fold change.

We used sparse partial least squares discriminant analysis (sPLS–DA) to globally identify the most discriminative transcripts that separated the sample groups, which were classified as components 1–3 (**Figure 1B**) (77). Component 1 (47%) distinguished tTregs from Tconvs regardless of costimulation (**Figure 1B**). Component 2 (13%) discriminated TNFR2-costimulated tTregs from CD3(/CD28)-stimulated tTregs and did the same, albeit more modestly, for Tconvs (**Figure 1B, left panel**). Component 3 (4%) highlighted a unique reaction of Tconvs to TNFR2 costimulation (**Figure 1B, right panel**). A representative heat map including transcripts with the highest contribution to components 1–3 showed that TNFR2-costimulated Tconvs and tTregs both expressed genes that set them apart from their CD3- or CD3/CD28-stimulated counterparts (**Figure 1C, Supplementary Table 1**). The transcriptomes of CD3- or CD3/CD28-stimulated tTregs and Tconvs clustered more closely together, especially in Tconvs, likely due to the convergence of CD3- and CD28 signaling pathways (78). The transcriptome analysis revealed in an unbiased manner that tTregs respond more strongly to TNFR2 costimulation at the level of gene expression as compared to Tconvs (**Figure 1D**). Gene Ontology (GO) enrichment analysis showed that processes related to mitosis were highly overrepresented in TNFR2-costimulated tTregs (**Figure 1E, left panel**). In TNFR2-costimulated Tconvs, GO and STRING network analyses indicated activation of TNFR family costimulatory pathways (CD70/TNFSF7, TNFSF9, RELB, MAPKs) and cytokine pathways (IL-12RB2, IL-21R, IL-23A) and inhibition of apoptosis (CFLAR/cFLIP, TNFAIP3) and inflammation (CASP-1 and −4, PYCARD, CARD16) (**Figure 1E, right panel; Figure 1F**). These data indicate that human Tconvs and tTregs both respond to TNFR2 costimulation, in a different manner, although the breadth of the transcriptomic changes is much larger in tTregs (16).

### 3.2 TNFR2 costimulation regulates metabolism in tTregs and Tconvs

Activated T cells rely on metabolic programs that support energy supply and biosynthesis needed for their rapid proliferation and effector functions (8, 9). Tconvs that are activated via the TCR/CD3 complex exhibit an mTOR-driven upregulation of glycolysis, which is promoted by CD28 costimulation (79, 80). Divergent results have been reported for Tregs (81-88), which may be explained by the complexity of the cell populations studied. For example, pTregs and tTregs are often not discriminated and cell populations often contain Tconv contamination. The impact of TNFR family members on cell metabolism has not been studied in any depth thus far. We recently reported that TNFR2 impacts T-cell metabolism. Specifically, we showed that upon TCR/CD3-mediated activation, human tTregs undergo an mTOR-driven glycolytic switch after TNFR2 costimulation, but not after CD28 costimulation, even though they enter cell division in both settings (16). As compared to TCR/CD3-activated glycolytic Tconvs, TNFR2-costimulated glycolytic tTregs show a net lactate retention and an increased flux of glucose-derived carbon into the tricarboxylic acid (TCA) cycle. Glycolysis proved essential to maintain FOXP3 expression and suppressive function in TNFR2-costimulated tTregs (16). Besides glucose, the amino acid glutamine is essential for T-cell proliferation and function (9). In Tconvs, glutamine uptake and catabolism is enhanced upon CD3/CD28 stimulation and is crucial for proliferation, differentiation and cytokine production (89-93). Glutamine is initially metabolized to glutamate and α-ketoglutarate, which can be further processed in the TCA cycle to support mitochondrial oxidative phosphorylation and the biosynthetic processes needed for cell proliferation.

We examined the metabolic processing of glucose and glutamine by TCR/CD3-activated tTregs and Tconvs in response to CD28 versus TNFR2 costimulation. We fed the cells [^13^C_6_]-glucose or [^13^C_5_]-glutamine and compared by mass spectrometry the processing of these nutrients into intermediates of the TCA cycle and nucleotide synthesis pathways. Tracing of [^13^C_6_]-glucose confirmed our previous finding (16) that TNFR2 costimulation promotes the influx of glucose-derived ^13^C into the TCA cycle in tTregs in particular (**Figure 2A**). Incorporation of ^13^C into citrate, α-ketoglutarate (both TCA cycle) or aspartate was increased in TNFR2-costimulated tTregs, whereas the increase was less pronounced in Tconvs (**Figure 2A**). Aspartate is a precursor for pyrimidine nucleotide synthesis, which can be analyzed by measuring UTP. Tracing of [^13^C_5_]-glutamine revealed that TNFR2 costimulation promoted both in Tconvs and tTregs incorporation of glutamine-derived ^13^C into α-ketoglutarate, aspartate and the nucleotide UTP (**Figure 2B**).

**Figure 2.**
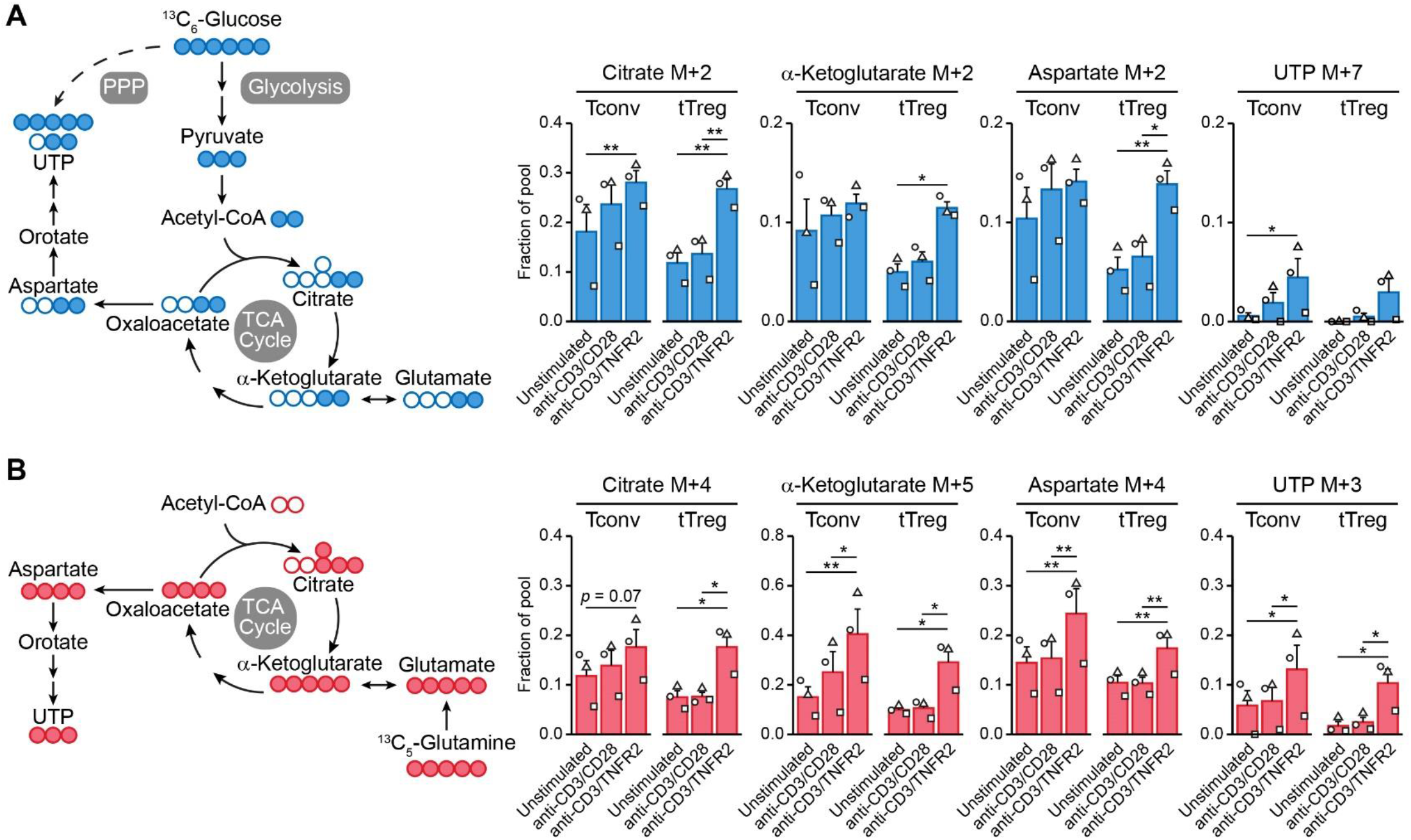
TNFR2 costimulation regulates Tconv and tTreg metabolism. Culture medium consisted of DMEM without glucose, pyruvate and glutamine (Gibco, Thermo Fisher), supplemented with additional non-essential amino acids (Gibco), 1 mM pyruvate (Gibco), 8% FCS and penicillin/streptomycin. To trace ^13^C-labeled nutrients, medium was supplemented with either 25 mM [^13^C_6_]-glucose and 4 mM [^12^C_5_]-glutamine (Cambridge Isotopes), or 25 mM [^12^C_6_]-glucose and 4 mM [^13^C_5_]-glutamine. In these media, pre-expanded human Tconvs and tTregs (5 × 10^5^) were restimulated in presence of IL-2, with or without agonistic mAbs against CD3/CD28 or CD3/TNFR2 (*n* = 3). After 24 h, cells were collected for metabolite analysis as described before (16), with minor changes. In short, liquid chromatography–mass spectrometry (LC–MS) analysis was performed on a Q-Exactive HF mass spectrometer (Thermo Scientific) coupled to a Vanquish autosampler and pump (Thermo Scientific). The flow rate was set at 100 µl/min. Quantification was based on peak area using TraceFinder software (Thermo Scientific). Peak areas were normalized based on total signal and isotopomer distributions were corrected for natural abundance. Tracing of [^13^C_6_]-glucose **(A)** and [^13^C_5_]-glutamine **(B)** is depicted in blue and red, respectively. Left: schematic diagrams of metabolic pathways following nutrient uptake, including glycolysis, the TCA cycle, the pentose phosphate pathway (PPP) and nucleotide synthesis. Right: quantifications of ^13^C-labeled (M+) citrate, α-ketoglutarate (both TCA cycle), aspartate and UTP (both nucleotide synthesis) as fractions of the pool. M+7 was shown for UTP in the [^13^C_6_]-glucose tracer experiment, as incorporated ^13^C originated from aspartate as well as PPP-derived ribose. Two-way ANOVA with Tukey’s post hoc test was used for statistical analysis (\**p* < 0.05, \*\**p* < 0.01). Data were log-transformed in case data were not normally distributed. Data are presented as mean ± SEM and data points are depicted as unique symbols per donor.

This is the first evidence that TNFR2 affects metabolism both in tTregs and Tconvs, with remodeling of glutamine metabolism as a commonality between both cell types and the enhancement of glycolysis and coupled TCA cycle events being unique for tTregs. Glutamine metabolism appears to promote nucleotide biosynthesis in both tTregs and Tconvs after TNFR2 costimulation, which can sustain cell proliferation. However, glutamine can also regulate effector differentiation: in CD4^+^ T cells it supports Th17 differentiation via glutamate and the antioxidant glutathione and impairs Th1 differentiation via α-ketoglutarate that acts as a cofactor for histone- and DNA demethylases (92). In this way, glutamine metabolism may support epigenetic reprogramming to maintain or direct tTreg and/or Tconv functionality under conditions of TNFR2 costimulation. This hypothesis remains to be proven, but we have already shown that glycolysis is essential to maintain FOXP3 expression and suppressive functions of tTregs after TNFR2 costimulation (16).

## 4 Discussion

Although TNF biology is very complex, TNF blockade is widely applied in disease and often successful. It has been well-established that TNFR1 and TNFR2 have distinct tissue distributions and signaling functions and that TNFR1 can be classified as pro-inflammatory, while TNFR2 is more anti-inflammatory. This realization fosters the approach of selective TNFR1- or TNFR2 agonism or antagonism in therapy of human diseases. Particularly, selective TNFR2 agonism to promote Treg responses and thus attenuate autoimmunity, inflammation and transplant rejection is highlighted in recent literature. Many data, including in vivo agonism of TNFR2 in mouse models argue that Tregs rather than Tconvs are the main responders to TNFR2 agonism. Even so, TNFR2 can also costimulate the proliferative response of Tconvs. We add here a side-by-side unbiased analysis of the response of human Tconvs and tTregs to TNFR2 costimulation at the transcriptomic level. This analysis highlights that both TCR/CD3-activated cell types have an inherent gene expression pattern that is discrepant between the cell types and does not change much upon CD28 or TNFR2 costimulation. We also observe a gene set that responds similarly to TNFR2 costimulation in tTregs and Tconvs, and a gene set that differentially responds between the cell types. These reactions are unique to TNFR2 costimulation and not shared with CD28 costimulation. These data suggest that the responder cells adopt a unique functionality upon TNFR2 costimulation that warrants further investigation. Furthermore, we highlight TNFR2 as a metabolic regulator—not only for tTregs, but also for Tconvs— as shown by a common upregulation of glutamine metabolism. It will also be of interest to investigate how this affects Tconv and tTreg responses.

Since tTregs are the major controllers of autoimmunity (72), we studied the response of tTregs to TNFR2 costimulation. The exact role of TNFR2 on pTreg responses remains to be elucidated. TNFR2 expression was not required for in vitro-generated, TGF-β-induced pTregs to suppress colitis in mice (94). Moreover, TNF blockade ameliorated EAE due to increased pTreg numbers, presumably by releasing an inhibitory effect of TNFR2 on TGF-β-induced FOXP3 expression (95). However, a contrasting study reports that TNF–TNFR2 signaling increases TGF-β-induced pTreg differentiation and suppressive function in vitro and in vivo (96). It remains to be unraveled whether the (metabolic) responses of human pTregs to TNFR2 costimulation are unique or more skewed towards the responses of Tconvs or tTregs.

Why Tregs as opposed to Tconvs predominate the response to TNFR2 agonists in vivo is not clear. One reason may be that TNFR2 expression is higher on Tregs than on Tconvs (16, 42, 97). By binding newly produced cell surface TNF, TNFR2^high^ Tregs may outcompete TNFR2^low^ Tconvs from responding to this ligand. By binding transmembrane TNF, Tregs may also prevent TNF from being shed and from activating TNFR1 on multiple cell types. It can be envisioned that Tregs in this way provide negative feedback to dampen inflammation (98). Certainly during priming in secondary lymphoid organs, proliferating Tregs and Tconvs are in close proximity. The precise contexts of TNFR2 costimulation remain to be resolved, but both CD4^+^ and CD8^+^ Tconvs (99, 100) as well as Tregs (101) can express transmembrane TNF. In a graft-versus-host disease model, TNF produced by Tconvs proved to be crucial for Treg responses (15). Activated Tconvs also produce IL-2 that Tregs need for their proliferation (76). Therefore, it is likely that Tconvs can invite their own suppression by Tregs via transmembrane TNF. Subsequent TNFR2-induced Treg proliferation may shift the balance—a numbers game—towards immunosuppression. In addition to Tconvs, monocytes (102) and tolerogenic monocyte-derived dendritic cells (103) can express transmembrane TNF, but how this impacts Treg or Tconv responses is not yet known.

After priming, Tregs relocate to non-lymphoid tissues, where they encounter Tconvs and other cell types. It has been shown that Tregs adapt their differentiation state in peripheral tissues to that of locally resident effector T cells, by responding to specific cytokines and gaining expression of lineage-determining transcription factors such as T-bet or GATA-3 in addition to FOXP3 (3). Since metabolic activity depends on nutrient availability in the environment, it is tempting to speculate that TNFR2 costimulation endows Tconvs and Tregs with a degree of metabolic flexibility that allows them to facilitate survival, replication and functionality in metabolically changing environments, such as sites of inflammation or cancer. TNFR2 is also involved in the function of myeloid-derived suppressor cells (104), mesenchymal stem cells (105) and multiple cell types of the central nervous system, including oligodendrocytes (106), their precursor cells (107) and microglia (108). The interesting commonality between these cell types and Tregs is that they possess immunomodulatory or tissue-regenerative properties. Similar to Tregs, these TNFR2-expressing cell types may exhibit a negative feedback mechanism to suppress the pathological effects of excessive TNF–TNFR1 signaling, e.g. in neurological disorders. Future studies are required to establish common TNFR2-induced (metabolic) responses of these cell types and how such responses may support their protective functions.

## 5 Data availability

Transcriptomics data are available in the GEO database under accession code GSE138604.

## 7 Author contributions

MM designed and performed cell stimulation and transcriptomics experiments and data analysis, prepared figures and wrote the manuscript, EAZ performed metabolomics and data analysis and prepared figures, TTNM performed transcriptomics data analysis and prepared figures, ES assisted with cell isolation, CRB and JB contributed to study design and writing the manuscript, SdK designed and performed cell stimulation and metabolomics experiments and contributed to study design and writing the manuscript.

## 9 Acknowledgements

We thank the staff of the flow cytometry facility and genomics facility at the Netherlands Cancer Institute and the flow cytometry facility at Leiden University Medical Center for their technical assistance. We also thank Lonneke Nouwen for support in transcriptomics analysis, Jeroen Jansen at the Utrecht Metabolism Expertise Centre for technical assistance, and Dr Sarantos Kostidis and Dr Martin Giera from the Center for Proteomics and Metabolomics at Leiden University Medical Center for insightful discussions.

